# Molecular Architecture of the human Citrate Synthase-Malate Dehydrogenase 2 metabolon

**DOI:** 10.1101/2025.10.13.681887

**Authors:** Angela J Kayll, Umanga Rupakheti, Renee St. John, Joseph J. Provost, Christopher E. Berndsen

## Abstract

Metabolons are transient biomolecular complexes that enhance the efficiency of metabolic pathways through substrate channeling. These complexes are difficult to study because of the transient nature, thus limiting our understanding of how they are formed and regulated. The citric acid cycle is proposed to contain many such complexes although few have been characterized structurally. Here, we provide direct structural evidence for the complex of human Citrate Synthase and human mitochondrial Malate Dehydrogenase 2, which is part of the larger proposed citric acid cycle metabolon. Our structural model supports previous crosslinking studies and suggests that hMDH2 can interact with each subunit of the hCS dimer, forming up to a hexameric complex. However, this complex appears transient as titration of hMDH2 into hCS in activity assays does not saturate. We further show that the interaction site with hCS is non-specific, as hCS could also stimulate oxaloacetate formation by cytosolic and plant MDH enzymes. This structural model will provide a base for understanding the structure and regulation of the broader citric acid cycle metabolon.

## INTRODUCTION

Cellular metabolism is a web of chemical reactions that must be regulated to produce complex life. Regulation and direction can take many forms, including protein post-translational modification, protein degradation, and coupling of enzyme reactions (Sweetlove & Fernie, 2018; Ciechanover, 2021; Provost *et al*., 2024; Martinez-Vaz *et al*., 2024). Malate dehydrogenase (MDH) is a conserved metabolic enzyme found in all branches of life that catalyzes the interconversion of malate and oxaloacetate in a NAD(P)(H) dependent fashion (Wolyniak *et al*., 2024; de Lorenzo *et al*., 2024). In most animals, MDH is found in the mitochondrion and the cytosol. In other branches of the tree of life, most notably plants, copies of MDH are also found in additional organelles (Baird *et al*., 2024; Wolyniak *et al*., 2024). In cellular metabolism, the function of MDH is best understood through thermodynamic coupling to CS, forming a metabolically linked enzyme pair that drives the tricarboxylic acid cycle forward (Omini *et al*., 2024). MDH catalyzes the “forward” reversible conversion of malate to OAA with the reduction of NAD^+^ to NADH. However, under standard conditions, the equilibrium of this reaction strongly favors malate, with a ΔG°’ of approximately +30 kJ/mol — making OAA formation thermodynamically unfavorable in isolation (Beeckmans & Kanarek, 1981; Vélot *et al*., 1997; Pettersson *et al*., 2000). The key to overcoming this barrier lies in the rapid consumption of OAA by CS, which irreversibly condenses OAA with acetyl-CoA to form citrate. This highly exergonic step effectively pulls the MDH reaction forward by depleting OAA, allowing continuous NADH production for oxidative phosphorylation. CS is also a rate-controlling enzyme of the TCA cycle, and its demand for OAA ensures that MDH remains functionally integrated into the cycle despite its thermodynamic constraint (Krebs, 1970; Elcock & McCammon, 1996; Vélot *et al*., 1997; Meyer *et al*., 2011; Wu & Minteer, 2019).

In 1973, Srere and colleagues demonstrated that the immobilization of malate dehydrogenase (MDH) and citrate synthase (CS) on a solid matrix enhanced the rate of oxaloacetate production, leading to the suggestion that MDH was in close proximity to CS within the mitochondrion, thereby resulting in a locally elevated concentration of oxaloacetate (Srere *et al*., 1973). Further work with glutaraldehyde crosslinking of MDH and CS resulted in enhanced reaction rates, suggesting the existence of a physical complex between the two proteins (Mattlasson *et al*., 1974). Subsequently, Halper and Srere isolated the MDH-CS complex in polyethylene glycol (PEG), exhibiting rate enhancement and specificity of porcine mitochondrial CS for mitochondrial MDH (Halper & Srere, 1977). In 1985, malate dehydrogenase was isolated as part of a larger complex with several other enzymes from the tricarboxylic acid (TCA) cycle, including citrate synthase (Robinson & Srere, 1985). The development of models depicting the MDH-CS complex, often involving other enzymes (Elcock & McCammon, 1996; Vélot *et al*., 1997), was subsequently followed by the investigation of fused MDH-CS proteins, which demonstrated enhanced activity in comparison to the free enzymes (Pettersson *et al*., 2000; Elcock *et al*., 1997). Several studies have shown a direct interaction between MDH and CS through cross-linking and mass spectrometry, leading to proposals for how these proteins interact (Wu & Minteer, 2015; Bulutoglu *et al*., 2016; Liu *et al*., 2017; Wu & Minteer, 2019). Moreover, this complex appears to be evolutionarily conserved, as MDH-CS complexes have been identified in proteins from pigs, mice, humans, yeast, and Bacillus subtilis (Shatalin *et al*., 1999; Meyer *et al*., 2011; Bulutoglu *et al*., 2016; Jung & Mack, 2018). The affinity between the two proteins is in the micromolar range, and the association is influenced by metabolites (Omini *et al*., 2021). However, observations of a complex revealing the relative orientation of the two enzymes and the stoichiometry of the complex remain absent. This is likely attributable to the transient nature of the complex.

Given the support for the MDH-CS complex over the past five decades, one might question the necessity of continuing to investigate the structure of this complex. Currently, it remains uncertain (1) how MDH facilitates the transfer of oxaloacetate to CS, potentially via a direct tunnel between the two enzymes, and (2) how this complex could be subjected to regulation (Pareek *et al*., 2021). Both MDH and CS form complexes with numerous other enzymes (Meyer *et al*., 2011; Omini *et al*., 2024), which may interfere with the formation of citrate or other metabolites; thus, a regulatory mechanism governing complex formation is required, contingent upon cellular status. Prior studies have presented conflicting evidence concerning the role of metabolites, such as ATP, in enhancing complex formation (Omini *et al*., 2021). Furthermore, it appears that MDH and CS co-diffuse at elevated rates of malate, a behavior that excess ATP moderates (Wu *et al*., 2015). The molecular basis underlying these observations remains unclear.

Considering the unresolved questions about the structure of the MDH-CS complex, we aimed to describe the solution behavior of these two proteins, including their potential interactions and the interaction site. Using small-angle X-ray scattering (SAXS), we characterized a crosslinked, hexameric complex of two human MDH2 molecules and one human CS. We also observed tetrameric forms of the complex, which could accommodate other binding partners. The interaction sites between the two proteins align with conserved surfaces on both proteins and seem to connect the active sites. Interestingly, we also found that we could not saturate CS with MDH in activity assays, suggesting that the hexamer we observe is a transient association. Additionally, we found that cytosolic MDH enzymes from plants and humans can function with human CS, while an MDH enzyme with a different surface charge density could not. These data support a structural model of the MDH-CS complex where MDH and CS bind through a charged surface, allowing the transfer of OAA from MDH to CS, followed by the dissociation of the complex.

## METHODS

### Purification of hMDH2 and hCS

Proteins with a hexahistidine tag were expressed in *E. coli* cells. The bacterial cell culture was streaked on an LB agar plate with antibiotic overnight in a 37ºC incubator. A single colony from a freshly streaked plate was placed into 100 mL of 2XYT media, shaking at 37ºC for 24 hours. 750 μL of the overnight culture was combined with 0.75 L of 2XYT media and 250 μL of 34 mg/mL Kanamycin. The solution was incubated with shaking at 37 ºC until the culture density reached OD_600_ 0.5-0.6. Once this was achieved, the temperature was decreased to 22 °C, 0.1 g of IPTG was added, and the culture was incubated overnight with shaking. To harvest the bacterial cells, the media was centrifuged with a Beckman Coulter Avanti J-26s XPI at 3,500 x g for 20 minutes at 4 ºC.

The proteins were purified using a multi-step chromatography method on an AKTA Start FPLC. We resuspended the hMDH2 or hCS bacterial cell pellets in 40 mL of 50 mM Tris-Cl, 100 mM NaCl, 1 mM Imidazole, and 0.1 mM EDTA, along with a Pierce protease inhibitor tablet (Fisher catalog number PIA32965) and 0.05 g of benzamidine. The pellet was warmed in a room-temperature water bath and vortexed until the solution was homogenous. Then, we sonicated the solution with a microtip at 60% power for three one-minute cycles 15 seconds on and 10 seconds off. We then centrifuged the sample in a Beckman Coulter Avanti J-26s XPI with a JA 25.50 rotor at 17,148 rcf for 30 minutes at 4ºC.

Once the pellet was prepared, we transferred it to an AKTA Start FPLC. The sample was initially purified with a HisTrap HP 1 mL Ni column. The lysis and wash buffer contained 50 mM Tris-Cl, 300 mM NaCl, 10 mM Imidazole, and 0.1 mM EDTA and the elution buffer contained 50 mM Tris-Cl, 50 mM NaCl, 300 mM Imidazole, and 0.1 mM EDTA. After loading the soluble lysate, the column was washed with 10.0 column volumes of wash buffer. The elution gradient was a linear gradient to 5% elution buffer in 10 column volumes, then to 40% elution buffer in 20 column volumes, and then up to 100% elution buffer in 10 column volumes. Afterward, we observed the fractions for a UV absorbance peak and confirmed the presence of protein in the sample using SDS-PAGE. Samples with protein of the appropriate molecular weight were centrifuged in 15 mL concentrator tubes with a MWCO of 3.5 kDa at 4 ºC until the sample volume was approximately 1.5 mL.

Following concentration, we further purified the proteins with a Sephacryl 16/60 S100 size exclusion column for hMDH2 and a Sephacryl 16/60 S200 size exclusion column for hCS in 50 mM Tris-Cl, 100 mM NaCl, and 0.1 mM EDTA. Afterward, we confirmed the presence of hMDH2 or hCS in the sample using SDS-PAGE. Samples with protein present, based on a band at the appropriate molecular weight, were centrifuged in 15 mL concentrator at 4ºC until the sample volume was approximately 1.0 mL. Protein concentration was determined using the extinction coefficients of 8,940 M^-1^ cm^-1^ for hMDH2 and 76,320 M^-1^ cm^-1^ for hCS.

### Cross-linking of hMDH2 and hCS

200 μL of 20 mM MES pH 6.4, 900 μL 1.3 mg mL^-1^ hCS, 300 μL 8.1 mg mL^-1^ hMDH2, and 1100 μL nanopure H_2_O were combined and incubated at room temperature for 15 minutes. 300 μL of 0.5% glutaraldehyde stock was combined with the protein solution and rapidly vortexed. The solution was incubated at room temperature for 30 minutes. We added 300 μL of 1 M TRIS base pH 8 to quench excess glutaraldehyde and then incubated the sample at room temperature for approximately 5 minutes. We then purified the sample with an AKTA Start FPLC and a Sephacryl 16/60 S200 size exclusion column in 50 mM Tris-Cl, pH 7.5, 100 mM NaCl, and 0.1 mM EDTA. Fractions containing crosslinked protein were identified with SDS-PAGE and concentrated in 15 mL concentrator tubes at 4ºC.

### Analytical SEC

We injected 50 uL of crosslinked MDH-CS onto a Wyatt 5 µm 300Å 4.6 mm ID SEC analytical column equilibrated in 20 mM sodium phosphate and 50 mM NaCl buffer at pH 8.0. The flow rate was 0.4 mL/min. Molecular weights shown in the figure were calculated based on the elution times of proteins in the Cytiva high and low molecular weight kits (product numbers 28403841 and 28403842). Standards were also used to generate a standard curve to calculate the molecular weight of the crosslinked protein complex.

### Small Angle X-ray Scattering

For hCS, we used SEC-SAXS and HT-SAXS to describe the structure, while we used SEC-SAXS for the hMDH2-hCS crosslinked sample. hCS was diluted to a concentration ranging from 0.07 mg mL^-1^ to 2.0 mg mL^-1^ in size-exclusion buffer (50 mM Tris-Cl, 100 mM NaCl, and 0.1 mM EDTA, pH 7.5) and then placed into a 96-well plate. The samples were frozen at -80 ºC before shipment. Data were collected at SIBYLS beamline 12.3.1 at the Advanced Light Source, Lawrence Berkeley National Laboratory (Classen *et al*., 2013; Rosenberg *et al*., 2022). Before sample collection, the sample plate was spun at 3700 rev min^-1^ for 10 minutes. Samples were held at 10 ºC during collection. The exposure was 15 seconds, with frames collected every 0.3 seconds for 40 frames per sample. The incident light wavelength was 1.03□Å at a sample-to-detector distance of 2.1 m. This setup results in scattering vectors, *q*, ranging from 0.013 to 0.5 Å^−1^, where the scattering vector is defined as *q* = 4πsinθ/λ, and 2θ is the measured scattering angle.

For SEC-SAXS, samples were separated on a Shodex KW-803 column at a flow rate of 0.5 mL/min at 10 °C, and eluate was measured in-line with UV/vis absorbance at 280 nm, Multi-Angle X-ray Scattering (MALS), and SAXS. The column buffer was 50 mM sodium phosphate, pH 7, 100 mM NaCl, 2 mM DTT, and 2% glycerol. The incident light wavelength was 1.03 Å at a sample-to-detector distance of 2.1 m.

To process SEC-SAXS data, Chromixs was used to subtract buffer from the peaks and produce the initial data file (Panjkovich & Svergun, 2018; Manalastas-Cantos *et al*., 2021). We then used RAW to analyze the data and generate statistics for the processed SEC-SAXS and HT-SAXS data and FOXS to fit structural models to SAXS data (Schneidman-Duhovny *et al*., 2013; Hopkins, 2024). The conformation of the hCS model was explored using NRGTEN and the models fitted to the data using FOXS [NRGETN]. The SAXS data and models are deposited in the SASBDB under codes SASDX87 and SASDX97 for hCS and SASDXA7 for the hMDH2-hCS complex.

### Lightdock

A dimer of hCS and hMDH2 dimers as constructed in Lightdock using crosslinking information from previous studies (Jiménez-García *et al*., 2018; Roel-Touris *et al*., 2020). The dimer of dimers was converted to a trimer of dimers via a two-fold rotation around the CS subunit. The resulting hexamers were fitted to SAXS data using FOXS.

### Coupling of MDH and CS activity

Assays coupling hCS and MDH activity were performed in 50 mM sodium phosphate (pH 8.0), with 100 uM malate, and 100 uM NAD^+^. Assays were run with 0.3 μM hCS and 0.4 μM of hMDH2 or 0.3 μM of the crosslinked complex or 0.4 μM of hMDH2 alone. Assays were performed in duplicate simultaneously. Assays were performed in a 96-well plate, and data were collected using a BioTek Eon spectrophotometer at 340 nm. Data were collected every 4 to 6 seconds for 10 minutes. MDH activity alone was confirmed in assays with 100 μM OAA and NADH instead of malate, NAD, and acetyl-CoA. MDH inactivity was confirmed in assays with 100 μM malate and NAD+ with no citrate synthase or acetyl-CoA, following the same data collection parameters.

### Titration of hMDH2-hCS activity

Reactions contained 50 mM sodium phosphate buffer (pH 8), 10 mM acetyl-CoA, 5 mM malate, 3.9 mM NAD, and 2.5 µM hCS. The hMDH2 concentration was varied to achieve hMDH2:hCS molar ratios of 0:1, 0.25:1, 0.5:1, 1:1, 1.5:1, 2:1, and 3:1. The hCS concentration was held constant. Final reactions were prepared by combining 40 µL buffer, 1 µL malate, 4 µL acetyl-CoA, 3 µL hCS, and corresponding volumes of hMDH2 in a final volume of 250 µL. Assays were performed using a BMG LABTECH CLARIOstarPlus device with Smart Control app at 25°C. To initiate reactions, 6 µL of NAD was added to each well, followed immediately by 200 µL of reaction mixture. Control reactions without the enzyme were included to account for background absorbance changes. Activity was measured at 340 nm for 150 cycles with 15 flashes per cycle and a cycle time of 5 seconds. Initial rates were calculated from the linear portion of the data and the data analyzed and visualized in R.

## RESULTS

### Structure of hCS

While the crystal structures of hMDH2 and hCS and their ability to interact are known (Bulutoglu *et al*., 2016; Schlachter *et al*., 2019; Berndsen *et al*., 2025), how these proteins associate and combine to form a metabolon is less clear. Before we could describe the complex, we first needed to conform the structure of hCS. Citrate synthase has not been characterized in solution via SAXS before while the solution structure of hMDH2 has been recently described (Berndsen *et al*., 2025). Thus, we needed to ensure that the X-ray crystal structures matched the solution behavior of these proteins to develop an accurate model of the complex. We compared out SAXS data on hCS to known crystal structures finding that hCS appeared to form a dimer in solution but the conformation of CS in crystal structures did not fully match our solution data (Schlachter *et al*., 2019) (Supporting Data Figure 1). Because hCS is likely to undergo conformational changes (Bayer *et al*., 1981; Schlachter *et al*., 2019), we sampled additional conformations of CS using normal mode analysis and found a conformation that fitted both data sets with X^2^ of 0.7 and 1.6 (Figure 1) (Mailhot & Najmanovich, 2021). In this conformation, one subunit of CS is in the open conformation with the active site accessible, while the other is in the closed conformation, which has previously been associated with a substrate-bound conformation (Schlachter *et al*., 2019)(Figure 1).

**Figure 1.**
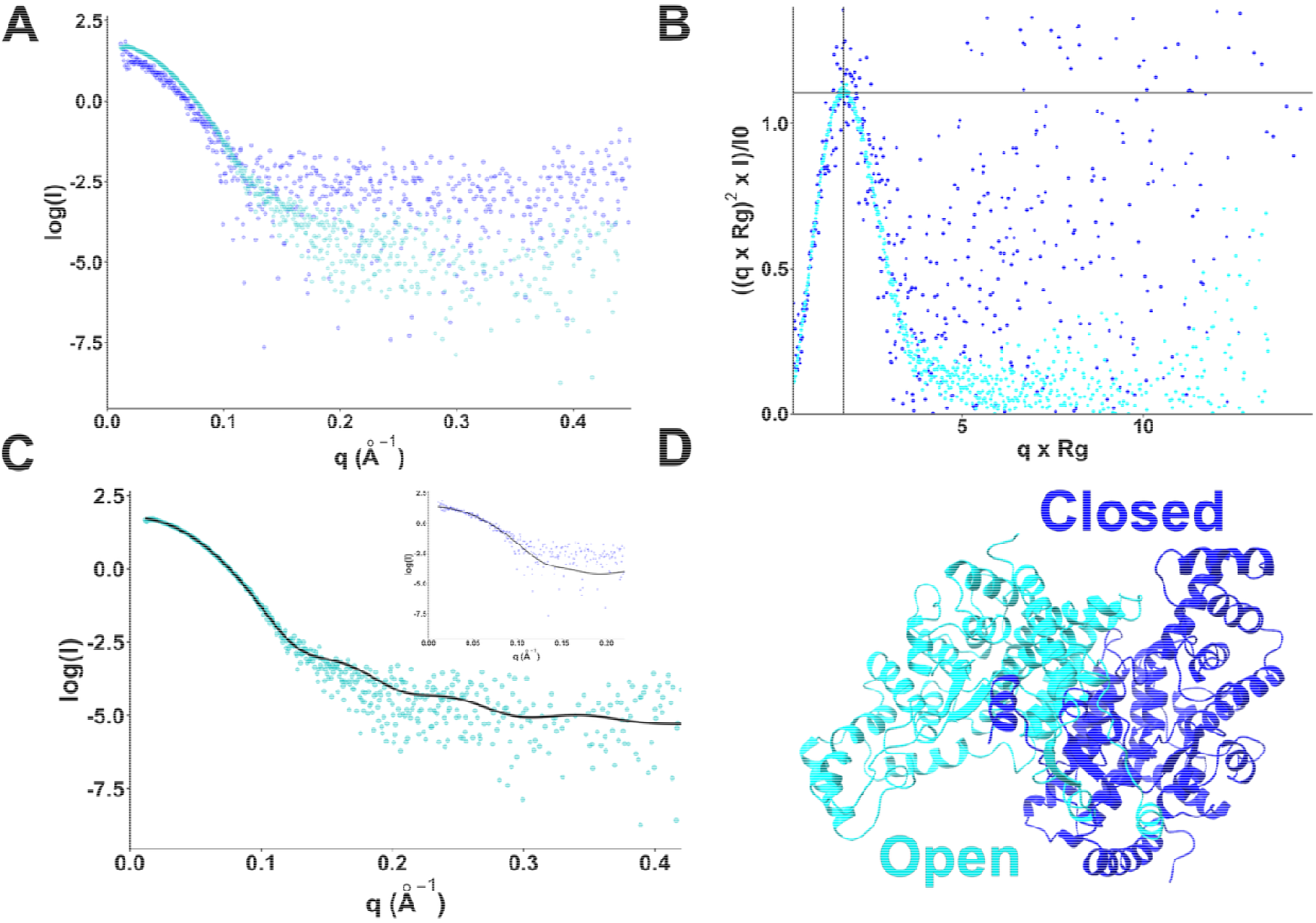
Small-angle X-ray scattering of hCS (A) log(I) vs. q plot for two data sets of hCS (B) Kratky plot of the data (C) Fitting of the modeled structure in (D) to SAXS data sets. The X^2^ value was 0.7 (main plot) and 1.6 (inset plot).

### Formation of the hMDH2 and hCS complex

Having more confidence in the solution structures hCS and hMDH2, we next worked to characterize the structure of the complex via SAXS. We initially tried to mix the proteins and purify the complex by size-exclusion chromatography. However, we failed to observe a prevalent and distinct complex of the two proteins (data not shown). Thus, we turned to glutaraldehyde crosslinking of the proteins to stabilize a potentially transient complex. We then purified the complex via preparative SEC followed by molecular weight determination by analytical SEC (Figure 1). We observed a single peak and found that the peak elution time was between the 150 and 220 kDa standard protein elution times, supporting the findings from the SDS-PAGE experiments. The calculated molecular weight was 187 kDa, suggesting a mixture of hexameric and tetrameric complexes. We separated factions on SDS-PAGE to see the resulting species. As shown in Figure 1, we observed a smear of bands consistent with a larger molecular weight complex of hCS and hMDH2. In fractions 5 and 6, this smear spanned from the 130 kDa marker to species larger than 170 kDa. Others have previously observed these higher molecular weight species using longer crosslinking reagents, suggesting that this complex is not an artifact of our system (Bulutoglu *et al*., 2016). These data suggest that hMDH2 and hCS can form a tetrameric or hexameric complex. Moreover, these data support a structural model where hCS has two possible interaction sites with hMDH2.

### Crosslinked hMDH2-hCS can form NADH and oxaloacetate

To confirm that the semi-purified, larger molecular weight protein complex observed in SDS-PAGE and analytical SEC was the hMDH2-hCS protein complex, and that crosslinking did not affect its function. The SEC purified fractions in Figure 1 appear to lack significant amounts of free hMDH2; thus, we felt confident that activity would be due to species in complex and not free enzyme. In NAD^+^ reduction assays, we compared the activity from solutions of hMDH2 and hCS with no glutaraldehyde crosslinking, crosslinked hMDH2 and hCS, and hMDH2 alone (Figure 1C). hMDH2 alone showed relatively little activity, suggesting it could not measurably reduce NAD+ in these assays. The cross-linked sample had activity levels within 2-fold of hMDH2 and hCS without crosslinking (Figure 1C). These data indicate that the crosslinked complex can generate NADH and oxaloacetate, the latter of which was likely used by hCS to produce citrate, although this was not directly measured. Previous studies on complexes linked by longer crosslinkers have shown similar activity (Shatalin *et al*., 1999; Bulutoglu *et al*., 2016). Therefore, any structural information obtained from this sample will probably reflect the complex’s catalytically active configuration.

### Structure of the hMDH2 and hCS Complex

We next collected HT-SAXS and SEC-SAXS data on the crosslinked MDH-CS. The HT-SAXS data showed evidence of higher molecular weight species, which we observed in the void region in the SEC-SAXS data. Thus, we used the SEC-SAXS data to model the configuration of the complex. The Kratky plot for the hMDH2-hCS complex, hMDH2, and hCS indicated compact, globular shapes (Figure 3 and STATS TABLE).

When comparing SAXS data from hMDH2 and hCS, to the hMDH2-hCS complex data, the higher values of radius of gyration (R_g_: hMDH2-hCS = 43.8 Å, hCS = 30.7 Å, hMDH2 = 28.1 Å), molecular weight (MW: hMDH2-hCS = 220.7 kDa, hCS = 96.1, hMDH2 = 68.8 kDa), and maximum length (D_max_: hMDH2-hCS = 150 Å, hCS = 99 Å, hMDH2 = 91 Å) observed for the complex suggest that the crosslinked sample are in an oligomeric state. If the sample contained a significant mixture of the free hMDH2 and hCS, we would expect these values to be between the values for isolated proteins. The data are more consistent with a mixture of tetramers and hexamers of hMDH2-hCS.

### Modeling the hMDH2 and hCS Complex

Initial attempts to use AlphaFold3 to generate a tetrameric or hexameric complex resulted in complexes that fitted poorly to the SAXS data (Abramson *et al*., 2024) (Figure 3C). We then used LightDock to create a tetrameric complex of hMDH2 and hCS with the restraint that R63 and R65 of hCS had to be adjacent to the interaction site to facilitate the transfer of oxaloacetate from hMDH2 to the active site of hCS (Jiménez-García *et al*., 2018). We created hexameric complexes from the tetramers by mirroring the MDH dimer across the hCS homodimer and fitting the models to the SAXS data. This configuration of hCS and hMDH2 in the model puts the oxaloacetate binding sites in hMDH2 and hCS from crystal structures 34 □ apart. The active site of hMDH2 aligns with the “basic pocket” surrounding the hCS active site, which is necessary for guiding oxaloacetate to the active site of hCS (Elcock & McCammon, 1996; Elcock *et al*., 1997; Wu & Minteer, 2015; Bulutoglu *et al*., 2016).

To support the stoichiometry suggested by the structural data, we titrated hMDH2 into hCS and measured the production of NADH in enzyme assays. Given that hMDH2 cannot produce significant amounts of NADH in the absence of hCS (Figure 1C), this assay setup is sensitive to the formation of the hMDH2-hCS complex. In fitting the data to a hyperbolic model, the data show a linear rise in activity until the hMDH2:hCS ratio is 1:1 followed by a gradual decrease in the steepness of the rise (Figure 4). However, we noticed a linear fit to the data was also visually reasonable. These data suggest that hCS did not saturate with hMDH2 under the assay conditions, suggesting a >1:1 ratio of hMDH2 to hCS, supporting the SAXS and SEC findings.

### Conservation around the interaction surface

We then used ConSurf to identify conserved surfaces of hMDH2 and hCS. To improve the results, we curated the sequences for each enzyme to only include known mitochondrial MDH and CS homologs and used the same species in each dataset (Supporting Figures 2 and 3). The interaction site within hCS shows greater conservation than other surfaces on CS (Figure 5). Moreover, this conserved patch includes the known interaction site and previously identified cross-linking sites (Bulutoglu *et al*., 2016). In hMDH2, the interaction is conserved and includes the active site of hMDH2. Thus, the atomic-level model of hMDH2-hCS is consistent with the SAXS data and is plausible based on biochemical and sequence conservation data.

**Figure 2.**
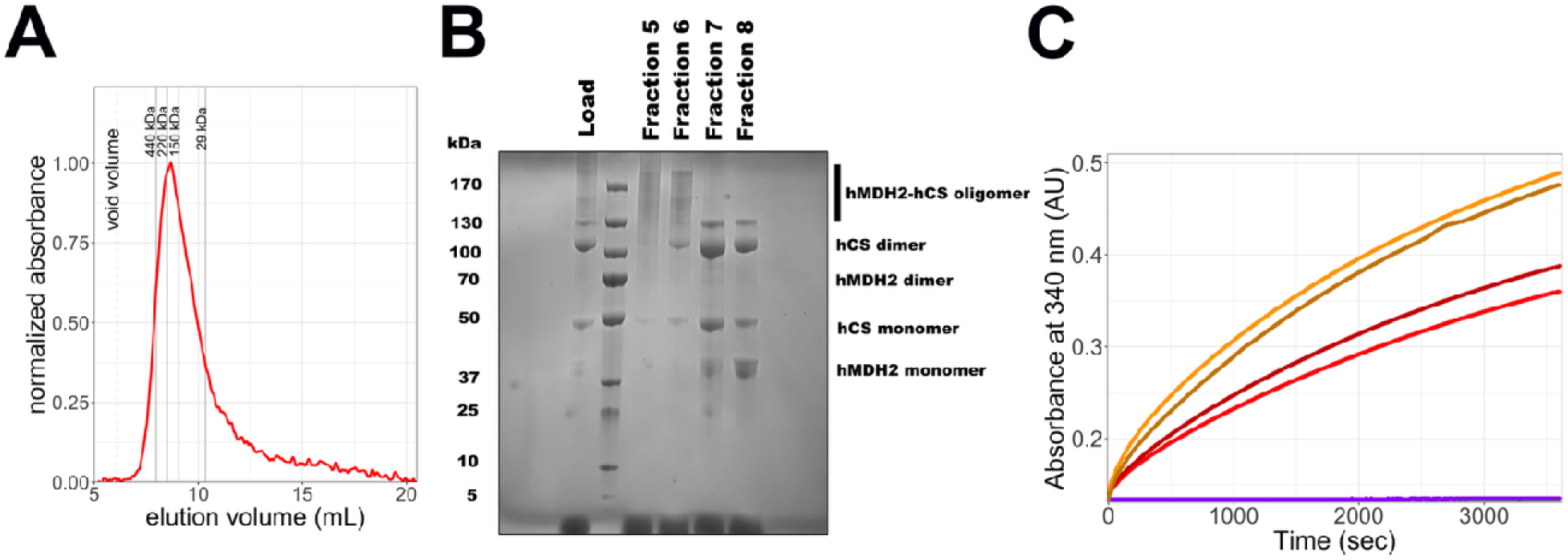
Purification of a crosslinked hMDH2-hCS complex (A) Size-exclusion chromatography of the crosslinked complex relative to the elution volumes of standard proteins (B) Gel of the main fractions across the peak in (A). (C) NAD+ reduction activity of the crosslinked complex (reds) relative to a 1:1 mixture of hMDH2 and hCS (orange) and hMDH2 alone (purple).

**Figure 3.**
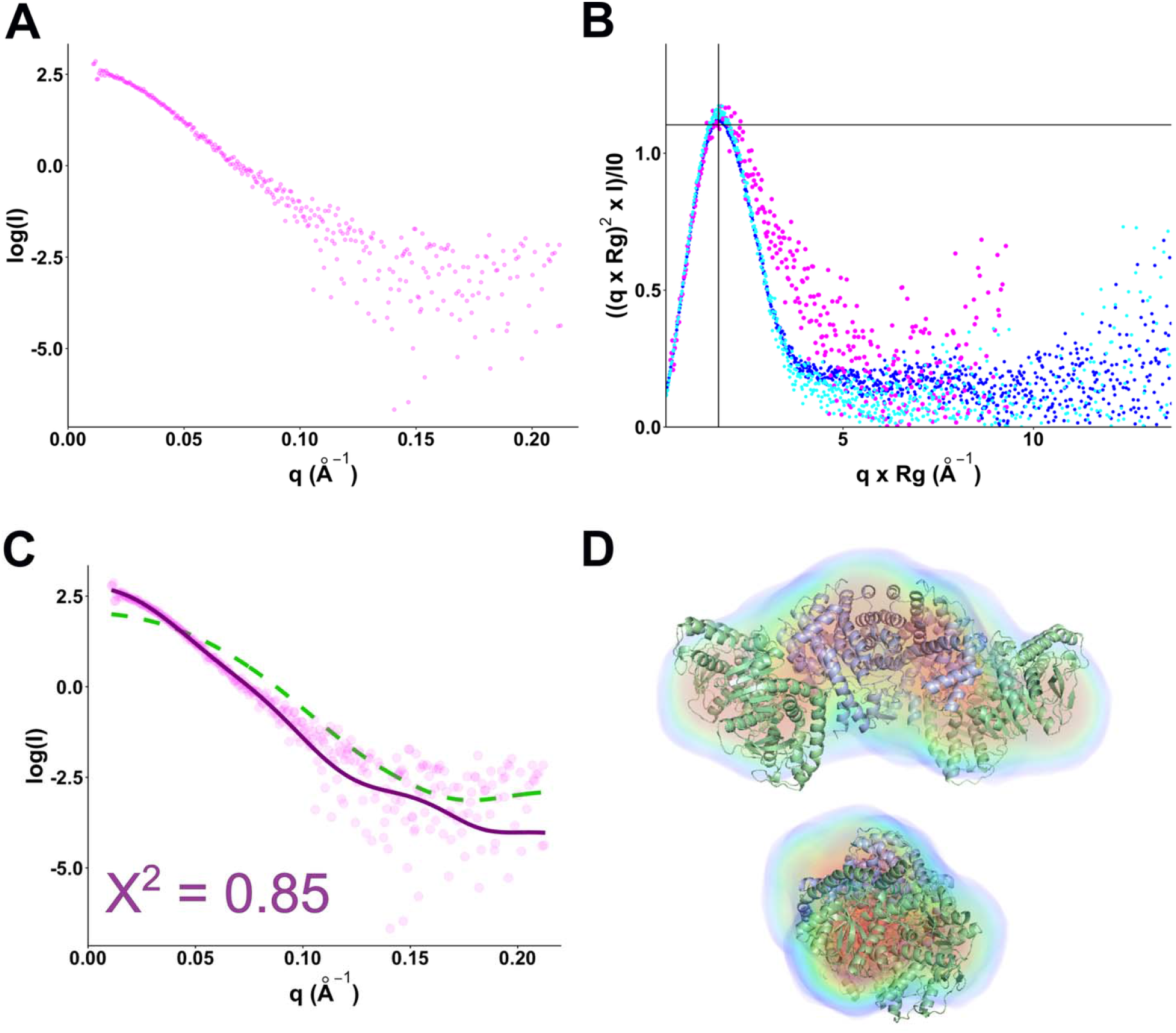
Small-angle X-ray data of the hMDH2-hCS complex (A) The log(I) vs. q plot of the trapped hMDH2-hCS complex (B) Comparison of hCS data from Figure XXX (blue/cyan) to the complex showing the differences in shape. (C) FOXS fitting of the AlphaFold3 model of MDH-CS hexamer (green, X^2^ = 2.3) compared to the final best fitting model derived from Lightdock fitting (purple, X^2^ 0.85). (D) Electron density of the MDH-CS hexamer derived from DENSS aligned to the best fitting hexamer model from (C). The mean real space correlation was 0.85 ± 0.04 and the Fourier shell correlation resolution was 55.6 ± 7.3 Å.

### hCS complex binding site is not sequence-specific

Given our findings that the hMDH2-hCS complex appears transient, we wanted to test the specificity of the interaction. First, we measured the OAA-forming activity of hMDH1 in the presence of hCS, finding that it was stimulated by hCS (Figure 6). We then expanded the specificity experiment to include a non-mitochondrial plant MDHs: the cytosolic MDH1 (AtMDH1) and the chloroplastic, NADP-dependent MDH both from *Arabidopsis thaliana*. We found that hCS also stimulated AtMDH1 activity, but that the NADP-dependent MDH was not affected, even in the presence of NADP^+^. Sequence comparison of all four enzymes showed limited sequence conservation of the interaction surfaces near the active site and the C-terminal regions implicated in binding to hCS. Prediction of the electrostatic surfaces of each of the enzymes showed a larger patch of negative charge and less positive charge in the NADP-dependent MDH that was not seen in the hMDH2 or the cytosolic MDH proteins (Figure 6C). These data support previous data that the interaction between MDH and hCS is partly defined by charge complementarity at the binding surface (Elcock & McCammon, 1996; Elcock *et al*., 1997; Shatalin *et al*., 1999; Wu & Minteer, 2015; Bulutoglu *et al*., 2016).

## Discussion

Interactions between adjacent enzymes in metabolic pathways are known to enhance the flow of metabolites and, in the case of hMDH2, help overcome unfavorable thermodynamic barriers (Sweetlove & Fernie, 2018; Omini *et al*., 2024). The structure and configuration of these complexes have been elusive, partly because of the inability to purify stable species. Previous work on hMDH2 and hCS has focused on the conditions that promote contact or identifying interaction sites (Halper & Srere, 1977; Bulutoglu *et al*., 2016; Omini *et al*., 2021). However, it was not clear how the two dimers fit together. Did the active sites of MDH pair with the hCS active sites? Was it a 2:1 MDH:CS stoichiometry? Thus, our work clarifies these previous studies to produce a structural model of the interaction consistent with the crosslinking studies and adds to our understanding of the complex. Here, we used glutaraldehyde crosslinking to stabilize a hexameric complex of hCS and hMDH2. This complex can form oxaloacetate *in vitro*, and similar complexes with masses of 170+ kDa have been seen by others (Bulutoglu *et al*., 2016). We further described the solution structure of hCS, finding that the crystal structures do not fit the data well due to asymmetry in the structure which is not fully seen in crystal structures (Schlachter *et al*., 2019). These data suggest a structural reaction mechanism for hCS that may involve cycling between conformations; however, this is not fully clear from the SAXS data presented. Modeling of the complex to SAXS data, guided by previous crosslinking studies, revealed that a hexameric model best fit the data (Figures 4 and 5) (Wu & Minteer, 2015; Bulutoglu *et al*., 2016). We then showed that the interaction surface is not species or family-specific, as hCS could promote oxaloacetate formation for cytosolic MDH enzymes from both humans and Arabidopsis. Our predicted interaction surface supports previous work showing that ionic strength was a strong regulator of complex activity, suggesting that the interaction site is charged and polar (Shatalin *et al*., 1999).

**Figure 4.**
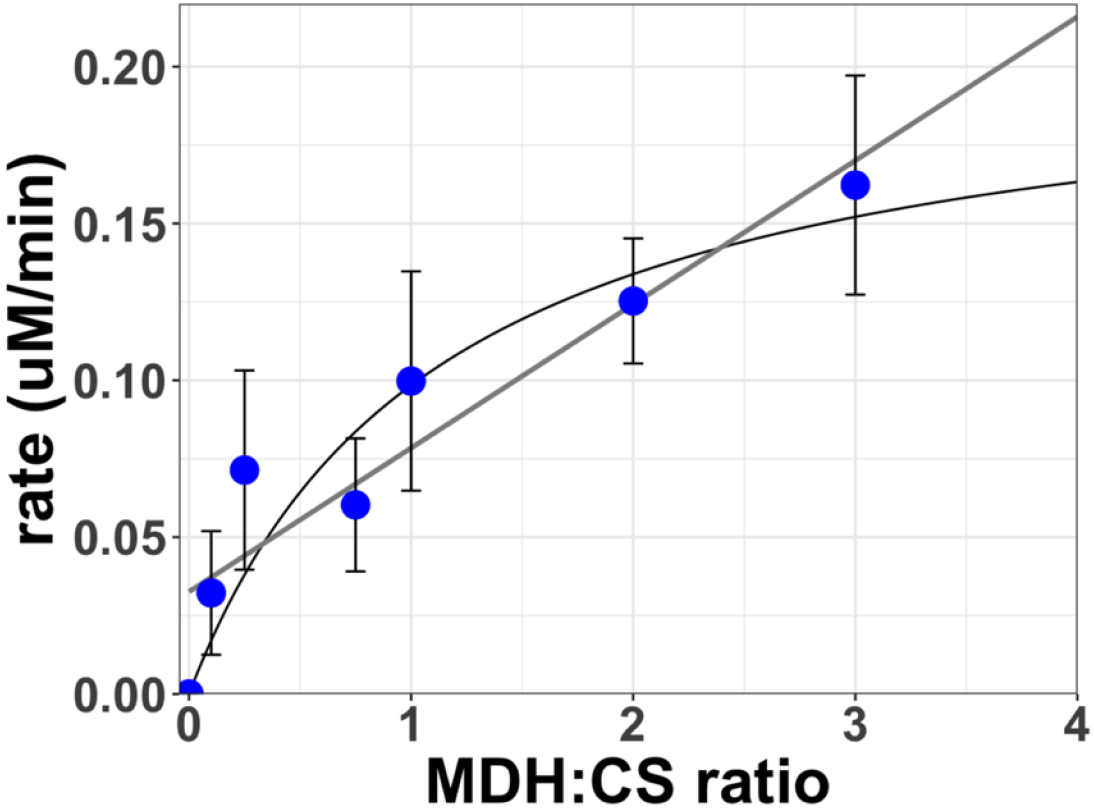
Titration of hMDH2 in activity assays. Blue dots are average values from at least three independent assays with error bars showing the standard deviation. The hyperbolic and linear fitting lines are shown, and the R^2^ values were 0.86 for the hyperbolic fit and 0.66 for the linear fit.

**Figure XX.**
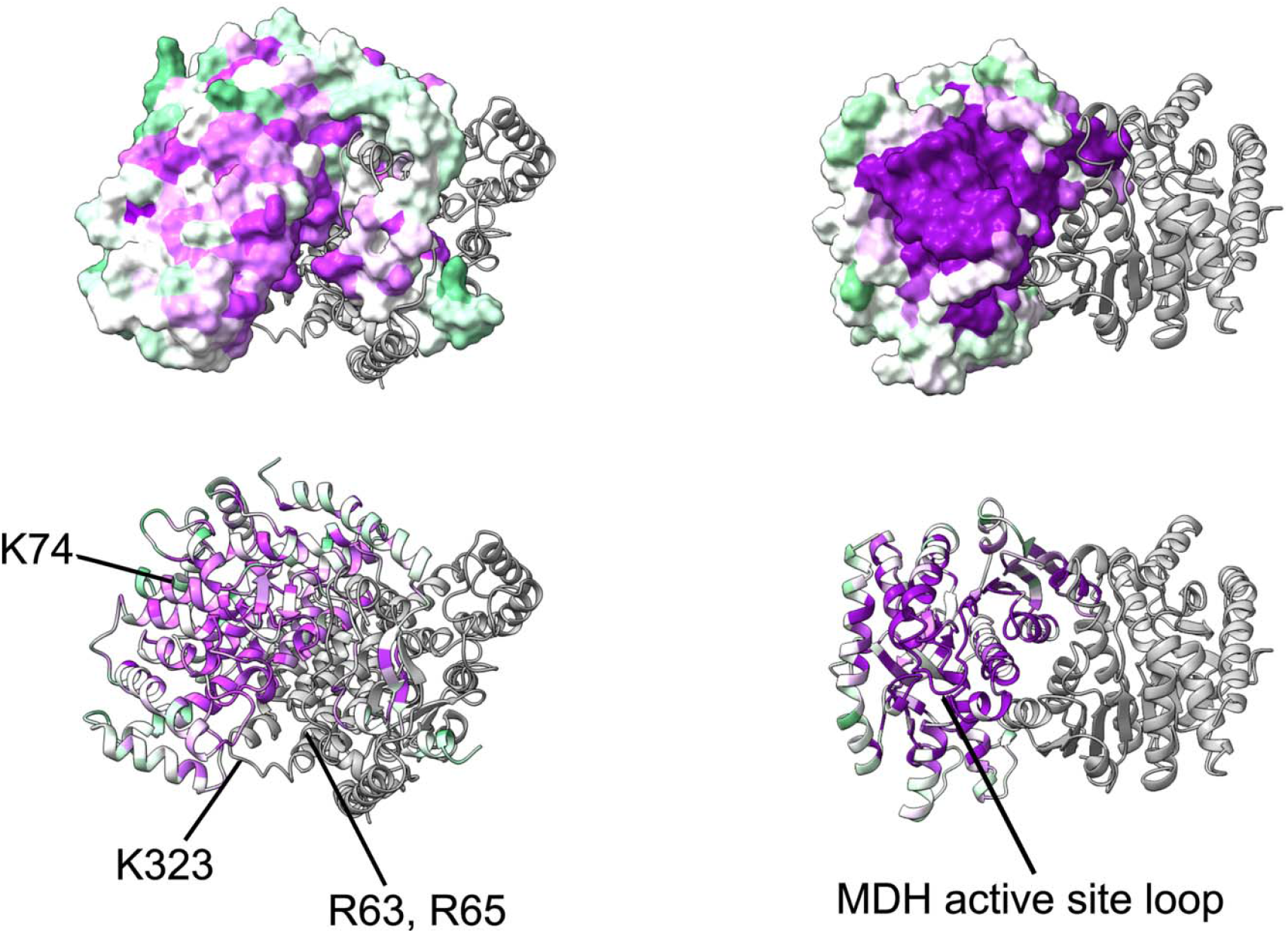
Conservation of the complex interaction surface for hCS (left) and hMDH2 (right). Identical amino acids in ConSurf analysis are indicated with purple coloring. Green shades indicate regions with 60-90% conservation, while white areas have less than 50% sequence conservation. Known interaction site side chains within hCS from previous work are indicated (Bulutoglu *et al*., 2016).

**Figure 6.**
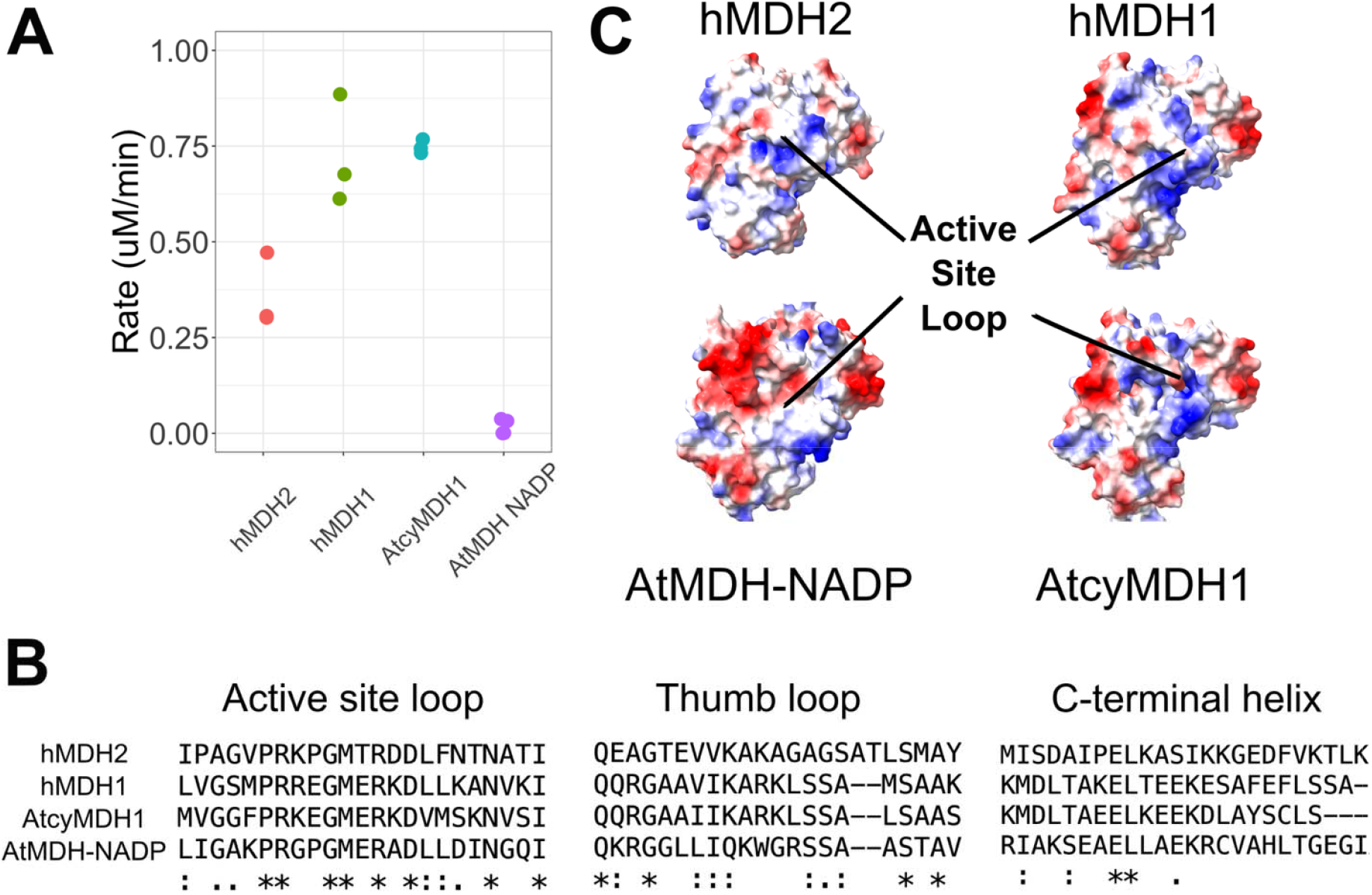
Interaction between hCS is based on surface charge – (A) assays of several MDH enzymes with hCS. Rate values from three separate assays are shown. (B) Conservation of the sequences of the active site loops and C-terminal helix in the hCS interaction site from Clustal Omega. (C) Surface charge of hCS site in the MDH enzymes used in (A). Red indicates negatively charged amino acid side chains, while blue colors indicate positively charged side chains.

From the structural model, the active sites of MDH and CS appear to “pair off” with a single MDH active site within the dimer interacting with a single active site within the CS dimer. This configuration maximizes the chances that OAA dissociating from MDH is proximal to the entry path of the CS active site (Bulutoglu *et al*., 2016; Schlachter *et al*., 2019). Moreover, this configuration is consistent with the reported reciprocal or half-the-sites behavior of the MDH dimer reported in the literature (Harada & Wolfe, 1968; Berndsen & Bell, 2024). In the reciprocal mechanism, only one active site in the MDH dimer is active at a time. In addition, CS is known to undergo significant structural rearrangements during the catalytic cycle near the interface with MDH (Bayer *et al*., 1981; Schlachter *et al*., 2019). Our SAXS data on hCS appear to further support this, as a model that was in a half-open/half-closed state appeared to fit the data better than the crystal structures. The structural rearrangements in both proteins are not blocked by interaction, as we and others show that crosslinked MDH-CS is an active complex (Bulutoglu *et al*., 2016; Shatalin *et al*., 1999). Thus, our proposed arrangement of hMDH2 and hCS is consistent with the catalytic cycle of both proteins.

The complex that we describe differs from the configurations proposed based on mass spectrometry and bioinformatics (Vélot *et al*., 1997; Bulutoglu *et al*., 2016). Even AlphaFold3 predicts a distinct configuration from what best fits our SAXS data. However, our predicted model does satisfy the contacts suggested by MS and does orient the active sites more closely than some of these models (Bulutoglu *et al*., 2016). Further, we provide higher resolution information by observing the predominant species in a solution of purified crosslinked protein via small-angle X-ray scattering. Thus, we consider our model a refinement of these earlier models based on new data, more so than presenting an alternative complex.

While we observed a hexameric species in SAXS, we also observed tetrameric species in other experiments, suggesting that both configurations are possible and functional. The trimer of dimers configuration is likely due to the excess hMDH2 that we used in the crosslinking reactions. Titration experiments of hMDH2 into hCS monitoring NADH production showed that hCS did not fully saturate under our experimental conditions suggesting complex formation is more transient. The historic difficulty in purifying this complex without crosslinking along with the low interaction site specificity support a model where MDH enzymes are rapidly transferring OAA to CS. Our findings further support recent work from Omini and coworkers and suggests a rapidly exchanging complex in yeast (Omini *et al*., 2025). This rapidly exchanging system would allow faster regulation of the system and simplifies the interaction site for a wide range of partners. Thus, our hexameric model likely represents a snapshot of a highly dynamic system and the maximally loaded state.

Since multiple interaction interfaces between malate dehydrogenase and citrate synthase have been observed, the factors that contribute to the stabilization or selective formation of specific MDH–CS complexes in vivo remain open questions. Additionally, how post-translational modifications might influence the assembly, orientation, or regulation of these metabolons is also under investigation. hCS is known to be methylated at K366 (K395 before processing), decreasing activity, and this modification site is adjacent to the interface with hMDH2 (Małecki *et al*., 2017; Rhein *et al*., 2017). Several large-scale studies have identified acetylation, phosphorylation, and succinylation of amino acids near the interaction site in both hCS and hMDH2, which would neutralize or invert the charge at the interaction surface (Zhao *et al*., 2010; Prus *et al*., 2024). In light of data suggesting charge complementarity as key for the hMDH2 interaction with hCS, these modifications would likely disrupt complex formation and activity. Further work is needed to confirm whether the other interaction site of hCS partners overlaps with the one described here; however, MDH and Aspartate aminotransferase are antagonistic in activity assays, suggesting a common surface (Beeckmans & Kanarek, 1981; Fahien *et al*., 1988; Elcock *et al*., 1997).

## Acknowledgements

We thank Dr. Jonathan Monroe for the critical reading of this work before publication. This work was supported in part by National Science Foundation grants MCB RUI-2322867 and CHE REU-2150091. This work was conducted in part at the Advanced Light Source (ALS), a national user facility operated by Lawrence Berkeley National Laboratory on behalf of the Department of Energy, Office of Basic Energy Sciences, through the Integrated Diffraction Analysis Technologies (IDAT) program, supported by DOE Office of Biological and Environmental Research. Additional support came from the National Institutes of Health project ALS-ENABLE (P30 GM124169) and a High-End Instrumentation Grant (S10OD018483).

