## Supplemental File for "Molecular Architecture of the human Citrate Synthase-Malate Dehydrogenase 2 metabolon"

By

Angela J Kayll, Umanga Rupakheti, Renee St. John, Joseph J. Provost, Christopher E. Berndsen

Supporting Figure 1

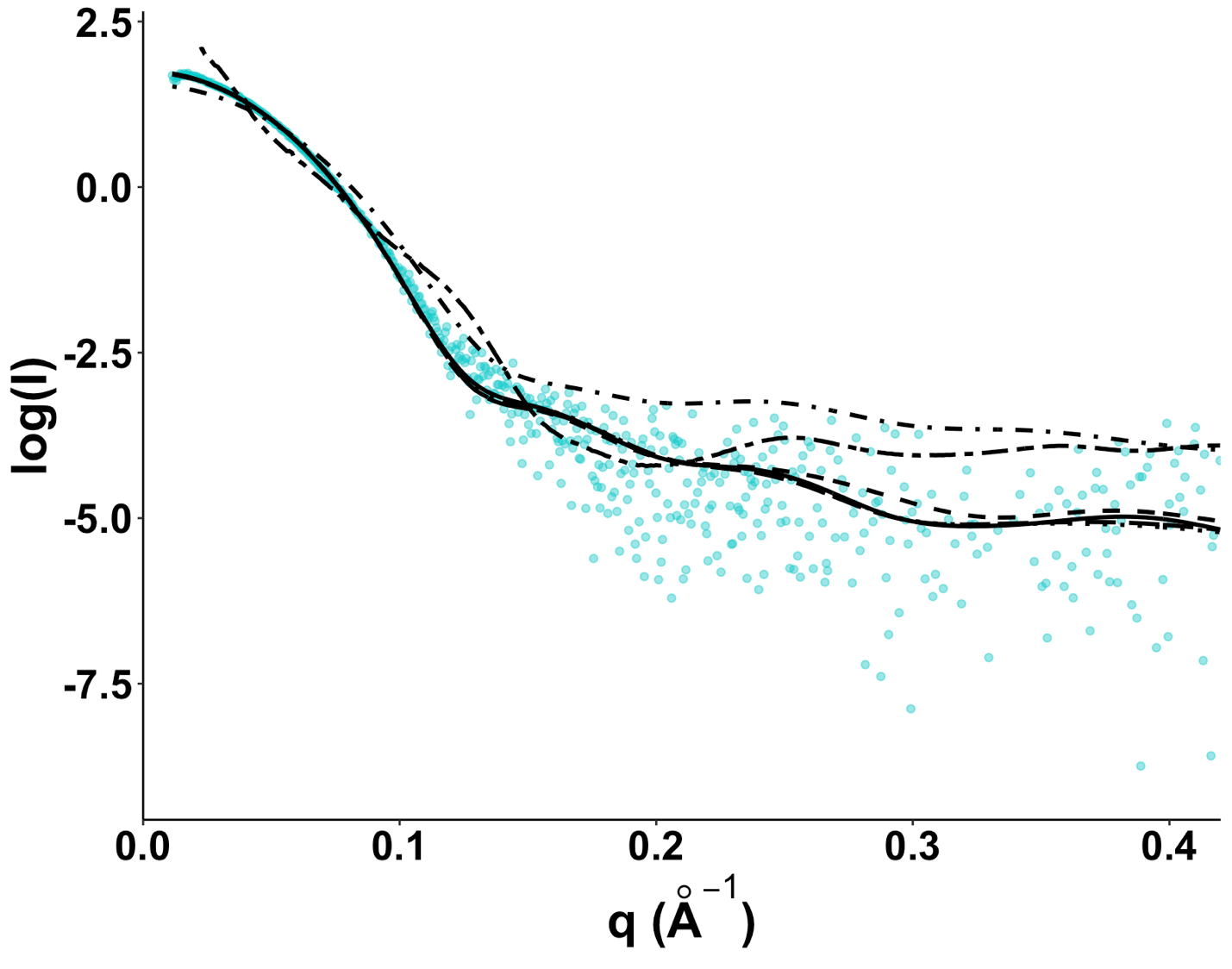

**Supporting Figure 1**: Fit of hCS SAXS data to crystal structures of citrate synthase. Predicted SAXS traces produced were fitted to the SAXS data in FOXS. PDB IDs used in fitting were 5UQU, 5UZP, 5UZQ, 6UZR, 6K5V, and 8GR8.

Supporting Figure 2

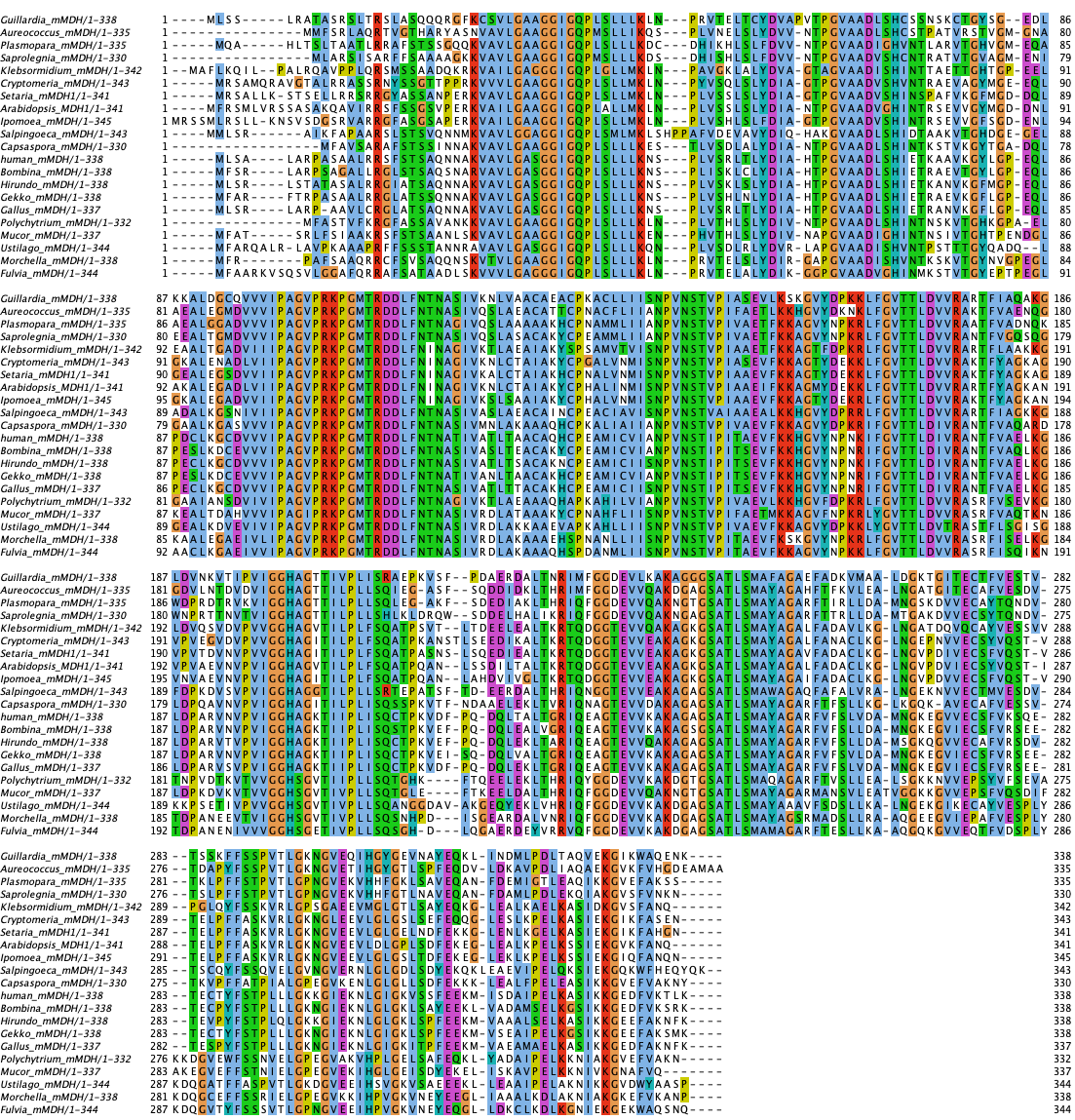

Supporting Figure 2: Alignment of Malate Dehydrogenase 2 homologs for ConSurf analysis.

Supporting Figure 3

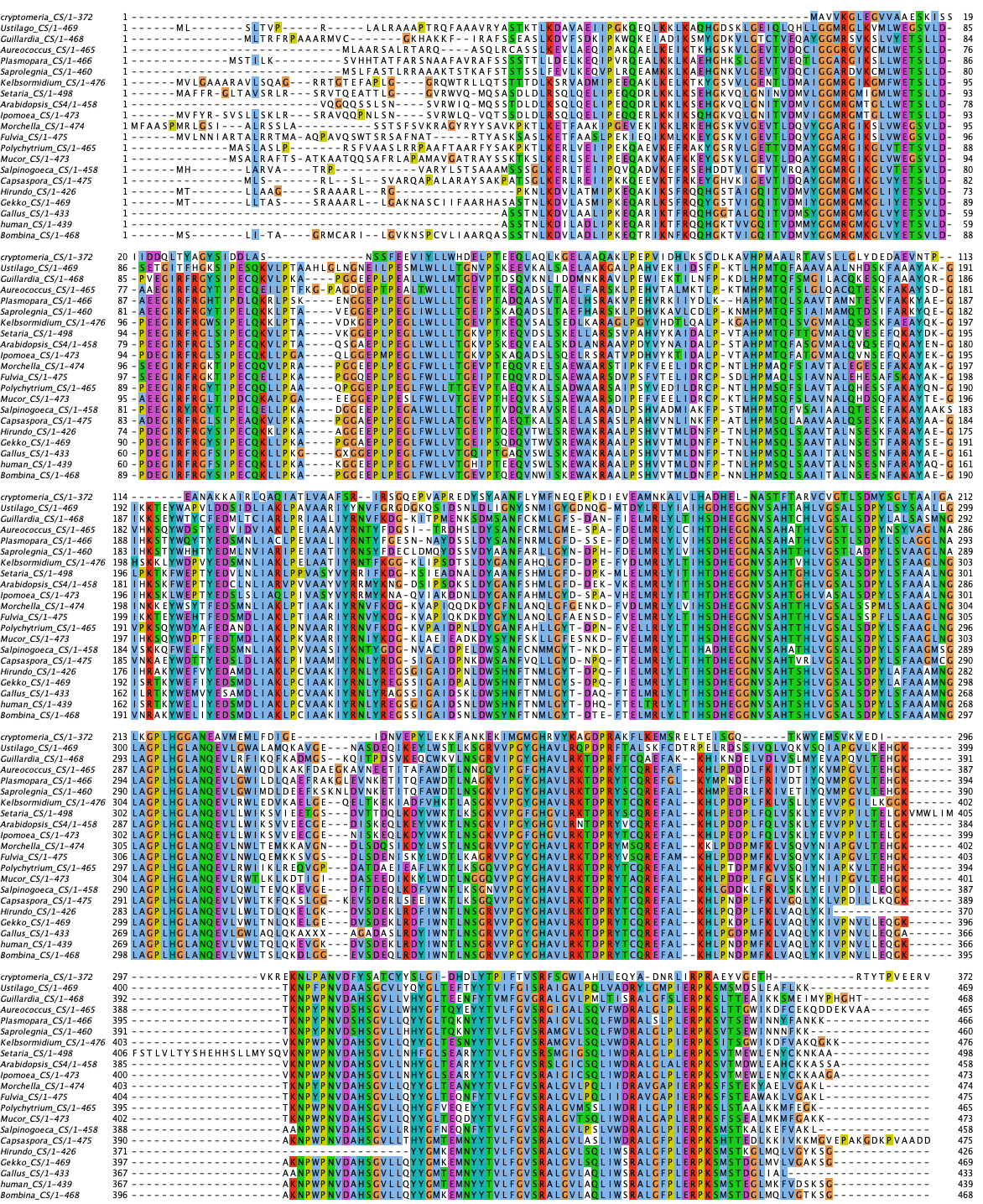

Supporting Figure 3: Alignment of Citrate Synthase homologs for ConSurf analysis.
